# Isolation of Elementary Nanofibrils of Cellulose from Non-Structural Plant Cells: Hydrothermal Processing as a Generalizable Route

**DOI:** 10.64898/2026.05.01.722164

**Authors:** Malak AbuZaid, M-Haidar A. Dali, Mohamed Hamid Salim, Vengatesan M. Rangaraj, Marjo Yliperttula, Fawzi Banat, Blaise L. Tardy

**Affiliations:** Department of Chemical and Petroleum Engineering, Khalifa University of Science and Technology, P.O. Box 127788, Abu Dhabi, United Arab Emirates; Food Security and Technology Center (FSTC), SAN Campus, Khalifa University, Abu Dhabi, United Arab Emirates; Faculty of Pharmacy, University of Helsinki, PO box 56, 00014 Helsingin Yliopisto, Finland

**Keywords:** nanocellulose, elementary fibrils, minimal processing, hazard-free, parenchymal cells, sustainable materials

## Abstract

The isolation of cellulose nanofibrils (CNFs), a promising precursor for sustainable and high-performance materials, has relied on chemically intensive, energy-demanding processes. As these processes were originally designed for the isolation of CNFs from wood, we herein show that the intrinsic ultrastructure of non-structural plant cells provides unique opportunities, namely direct access to loosely organized cellulose nanonetworks. We demonstrate that this loose nanofibrillar tissue can be transformed into CNFs with sizes down to elementary nanofibrils (∼4 nm) at high yields (reaching ∼32%) under exceptionally mild hydrothermal conditions. Three distinct plants were evaluated and the physicochemical properties of the obtained nanonetworks and corresponding CNFs were thoroughly studied, including the hydrodynamics of the resulting gels. Films prepared from the obtained CNFs showed similar performance to those obtained from conventionally isolated wood-based CNFs. Overall, this study demonstrates that CNFs can be obtained through low-intensity, hazard-free, processes from widely available biomass. Thus, this approach offers a unique shift in the range of opportunities to produce CNFs facilitating the integration of their use into the food supply chain, biomedical applications, and other regulatory-constrained applications.

## INTRODUCTION

Cellulose is the backbone of the global bioeconomy, either as a bioproduct source from the forest sector, in the form of agricultural products, or as animal feed. Yearly, over 40 billion tons of cellulose are biosynthesized by nature (Hon and N.S 1994), principally in the form of compact bundles of elementary nanofibrils with a diameter of ca. 4 nm (Zhu et al. 2015; Beaumont et al. 2021). The efficient isolation of these cellulose nanofibers (CNF) has thereafter been one of the main goals associated with the next generation of sustainable materials, which combine high performance, versatile engineerability, and ecosystem integration (Li et al. 2021b). To date, the production of nanocelluloses from plants has been widely explored including annual crop residues, food losses (Thomas et al. 2018) and, most extensively, wood (Li et al. 2021b; Hamedi et al. 2025). Critically, all developed processes are derived from those originally designed for wood, the first plant from which nanocellulose was efficiently isolated (Gómez H. et al. 2016; Ma et al. 2022). These processes involve severe chemical pre-treatment steps and high-intensity mechanical post-treatment, regardless of the plant-biomass original ultrastructure or composition.

Fundamentally, the cell wall structure and composition of woody plants differ substantially from those of non-woody plants. The majority of wood cells’ functions fulfill a considerable load-bearing component, which result in densely lignified tissues (Pennells et al. 2020). Wood requires high severity alkali pre-treatments (e.g. Kraft Pulping at 150 □C, 7 bar, 15 wt.% reagents), followed by bleaching and high intensity mechanical processes, typically high-pressure homogenization to yield branched and bundled CNFs (Mattos et al. 2019). Milder processes have been developed for the isolation of individualized nanofibrils from non-woods; however, these always involve a chemical or biochemical pre-treatment (Li et al. 2020; Amoroso et al. 2022; Jančíková and Jablonský 2024; Ren et al. 2024; Fang et al. 2025). The cells of non-wood feedstocks, e.g., those from food wastes and losses, have thinner and less stratified cell walls, where the secondary wall of parenchymatous tissues of edible plants can even be absent (Wang et al. 2018). Furthermore, elementary fibrils of cellulose in soft plant tissues were reported to be separated by ∼20 nm in their cell walls (onion cell-wall (Ye et al. 2018)), compared with a few nanometers in wood (Fernandes et al.; Cosgrove 2014). They have low lignin contents, typically between 1 to 15% (Wang et al. 2018; Pennells et al. 2020; Gröndahl et al. 2021; Ciriminna et al. 2024), water contents that can exceed 90%, resulting in “soft” mechanics for the corresponding plant tissues (e.g., a compression strength of 0.64 MPa for carrot (Xu et al. 2015) compared to 60 MPa for birch wood (Al-musawi et al. 2024)). Collectively, these properties suggest that the cellulosic nanonetwork in non-wood plant tissues may be highly accessible and considerably easier to nanofibrillate than wood, thus, offering the potential to: (i) substantially decrease mechanical processing energy use and production costs, (ii) generalize the removal of chemical in pre-treatments, and (iii) eliminate or largely reduce safety concerns and strict regulations applied to traditional, chemically pre-treated nanocelluloses (More et al. 2021; Qin et al. 2024). The latter represent all three key bottlenecks associated with a wider-scale implementation of CNFs across all sectors, where their performance has been demonstrated repeatedly.

Herein, we hypothesize that non-wood plant tissues can exhibit dramatically lowered thresholds to nanofibril isolation as associated with their inherent ultrastructure, which extends beyond differences in compositional differences. To validate this hypothesis, three distinct plant tissues, namely carrots (*D. carota subsp. sativus*), watermelon (*C. lanatus*) and dates (*P. dactylifera*) were chosen for their low lignin content and inherently soft mechanics associated with the predominant storage functions of their cells. The tissues were analyzed for their intrinsic cellulosic nanonetwork as well as their potential for CNF isolation by minimal processing (**Figure 1c**). Hydrothermal treatments (∼80-98□) at atmospheric conditions (**Figure 1c_1,2_**) and mild shredding (**Figure 1c_3_**) uniquely enabled isolation of high-grade cellulose (**Figure 1a**), with substantial upsides compared to the state of the art in isolation processes (**Figure 1b**). Diffraction, infrared and UV-Vis spectroscopy analysis as well as atomic force and electron microscopy techniques revealed loose cellulose nanonetworks in tissue and high-grade CNFs from subsequent isolation, with few non-cellulosic residuals. For all three plants, a near-complete transformation to nanofibrils was observed, with gelation concentrations ranging from 0.68% to 2.0% and diameters comparable to those observed for elementary fibrils. Finally, the formulations obtained were trialed for simple nanopaper fabrication and benchmarked to those obtained from wood, showing comparable performance. This work showcases that high grade CNFs can be isolated at high yields (up to 32 wt%) through a process estimated to be at least 80% less severe to the state of the art, with substantially reduced associated costs. We expect those results to be highly generalizable across a wider range of plant tissues as the three cases evaluated herein encompass a wider range of plant types that are typically readily available as co-products of the food chain. Overall, this study provides an opportunity to overcome the current barriers behind a wide-scale implementation of CNFs as high-performance and sustainable building blocks.

**Figure 1.**
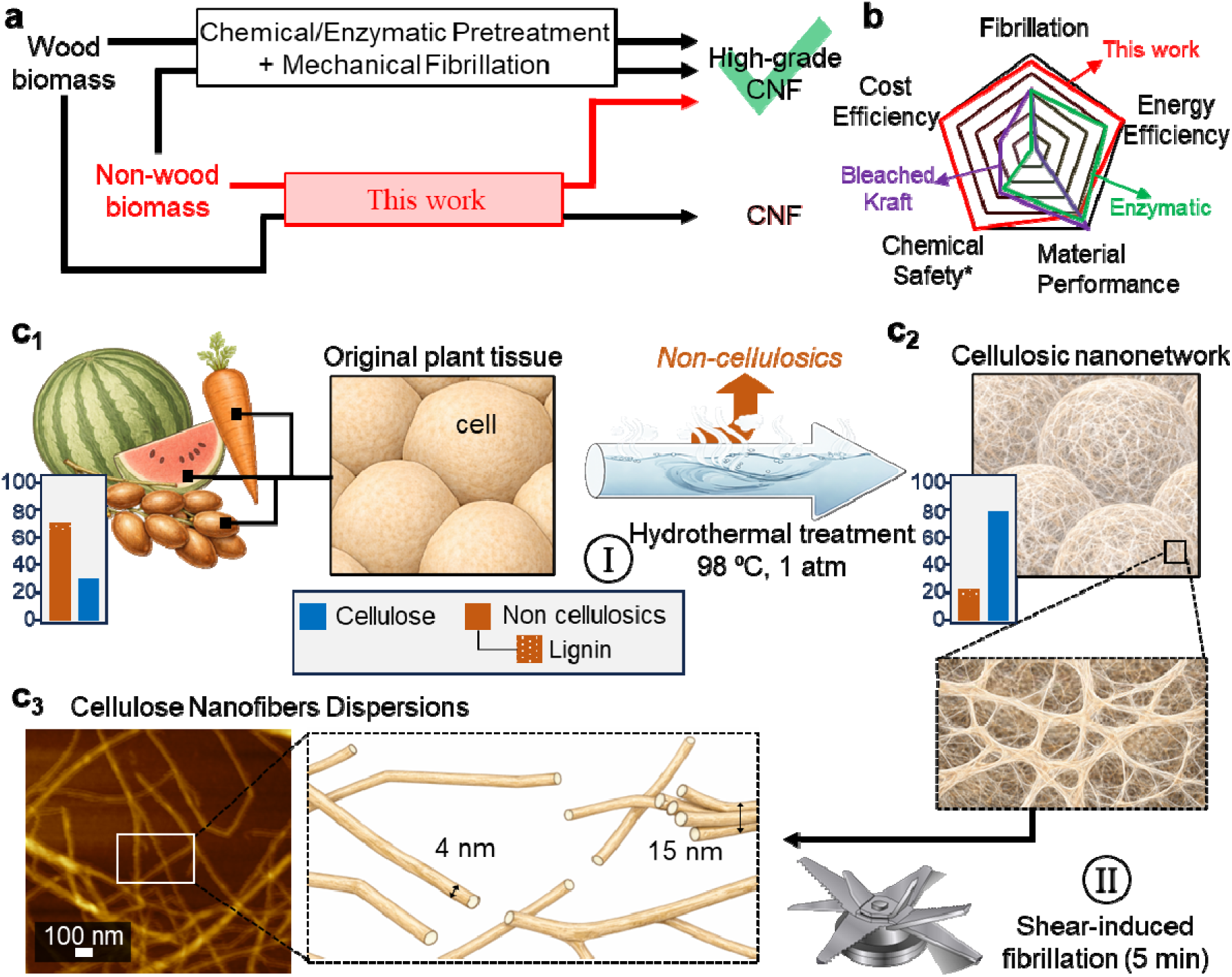
Comparison between conventional, wood-derived, CNFs isolation and the present ultrastructure-enabled route. (a) Schematic comparison of conventional wood-based CNF production (chemical/enzymatic pre-treatment followed by mechanical fibrillation), with the present approach using non-wood biomass to directly obtain high-grade CNFs. (b) Radar plot benchmarking of the proposed method against two wood-based routes (obtained as per method provided in the experimental section). *chemical safety herein refer to process safety rather than the use of the obtained CNFs across applications. Illustration of the proposed method; (c_1_) Hydrothermal pre-treatment (80-98°C (upper bound shown), 1 atm) removes non-cellulosic components and reveals loose cellulose nanonetworks, where (c_2_) the subsequent 5 minutes mild shear-induced fibrillation yields bundled and individualized nanofibrils down to elementary nanofibril sizes.

## RESULTS AND DISCUSSION

To test the hypothesis that some cellular ultrastructure would enable unprecedented nanofibril accessibility, we selected plant tissues with predominantly non-structural functions and low lignification. Date (D), carrot (C), and watermelon (W) inner tissues, as well as carrot and watermelon peels (CP and WM, periderm (Tian et al. 2024) and mesocarp, respectively) were initially evaluated for their inherent nanonetwork (**Figure 2a_1_, S2a_1_, S3a_1_**). The tissues were treated by hydrothermal processing under atmospheric conditions, 1 atm and ca. 98L. Following hydrothermal pre-treatment (with corresponding samples bearing an additional ‘Hydro’ subscript), a change in color occurred (**Figure 2a_2_, S2a_2_, S3a_2_**), along with a softened structure, as commonly observed. These alterations indicated substantial removal of weakly bound non-cellulosics such as hemicelluloses, polyphenols, and pectin, accompanied with partial pigment and water□soluble nutrient leaching such as flavonoids and carotenoids. A fibrous structure was readily visible by the naked eye in the treated date fruit and was further exposed upon freeze-drying using tert-butyl alcohol to prevent freeze-templating by ice crystals (Inoué and Osatake 1988; Govindharaj et al. 2025) (**Figure 2b**). Scanning electron microscopy (SEM) of the treated biomass highlighted thin cell walls and loose nanofibrils present in the resulting nanonetworks (**Figures 2c–e, S1, S2b, and S3b**), also corroborating with continuous entangled cellulosic nanonetworks showcased previously for hydrothermally treated carrots (Varanasi et al. 2018). Across the different selected tissue types, variations in cell size were observed, with cross-sectional sizes of the lumen ranging from approximately 16.7 µm for carrot periderm up to 318 µm for watermelon mesocarp. The largest cells were identified in watermelon, particularly within the mesocarp region. Such large and thin, cell wall thickness ∼250 nm, non-structural, i.e. parenchymal, cells are characteristic of tissues with high water content, where their primary function is the storage of water, soluble sugars, nutrients (Liu et al. 2024), and metabolites, rather than providing mechanical support. Consequently, in these tissues, pectin, cellulose, and hemicellulose are predominant, while lignin is nearly absent, unlike in structural or vascular regions (e.g. in the xylem) where lignification contributes to rigidity (Zhang et al. 2025). Distinct nanofiber networks were evident in all samples with wide gaps of over 20 nm between nanofibers, aligning with previous diffraction analysis (Ye et al. 2018). This confirmed that loose nanofibers were inherently present within the biomass, and suggested that the hydrothermal pre-treatment effectively removed non-cellulosic components, exposing the underlying cellulose nanonetworks.

**Figure 2.**
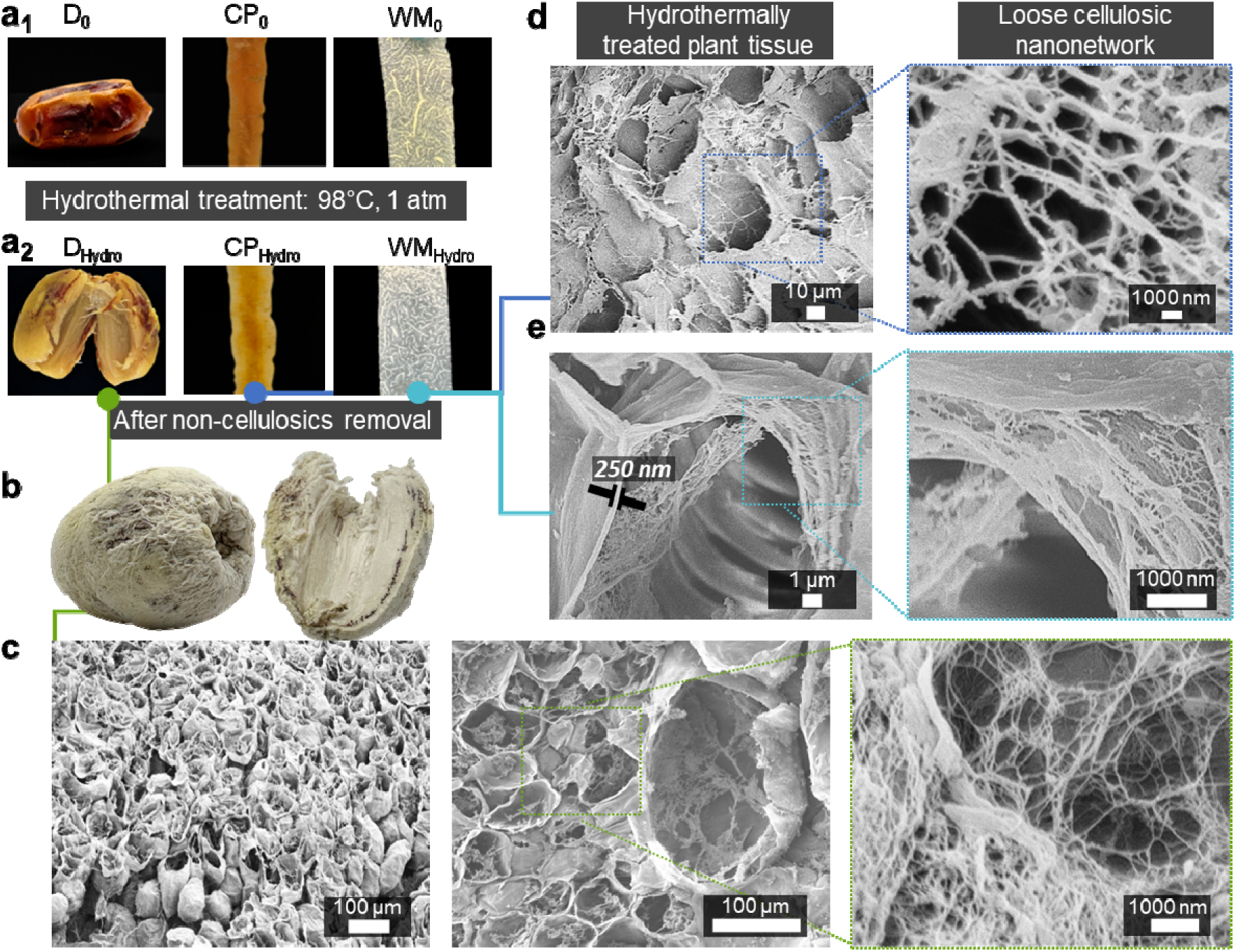
Ultrastructure of hydrothermally treated non-structural plant tissues. (a) Images of date (D), carrot periderm (CP), and watermelon mesocarp (WM) tissues (left to right, respectively) (a_1_) before (D_0_, CP_0_, WM_0_) and (a_2_) after hydrothermal pre-treatment (D_Hydro_, CP_Hydro_, WM_Hydro_). (b) Date after non-cellulosics removal and water removal by lyophilization. (c-e) Scanning electron microscopy (SEM) micrographs of hydrothermally treated (c) date (D_Hydro_), (d) carrot periderm (CP_Hydro_), and (e) watermelon mesocarp (WM_Hydro_).

Conventional pretreatments typically involving exposure to a strong acid or base at high temperature followed by bleaching previously demonstrated the isolation of clearly individualized nanocelluloses (i.e. as contrasting to nanonetworks) (Ren et al. 2024; Fang et al. 2025). Considering carrot, for example the mildest process that showcased individualized nanofibrils used a 2h treatment at 80L in water followed by peracetic acid bleaching at 90°C for 2 h followed by 10 passes of high-pressure microfluidizations (Amoroso et al. 2022). Hydrothermal processes coupled with intense mechanical processes (PFI mill refining and multi-pass high-pressure homogenization) led to exposed nanofibers, hinting at possible individualized nanofibers (Varanasi et al. 2018). Enzymatic pre-treatments have also been effective, though similar mechanical post-treatments are required (Fang et al. 2025). New solvent systems such as deep eutectic solvents (DES) have also been explored (Li et al. 2020; Jančíková and Jablonský 2024; Fang et al. 2025), for instance yielding okara-derived cellulose nanofibers with an average diameter of 27 nm following DES pre-treatment and high-pressure homogenization (Li et al. 2020). To date, the mildest processes reported for the isolation of individualized nanofibers of cellulose are for non-woods and can bypass the use of high pressure homogenization, though severe chemical or biochemical pre-treatments are still used (Dali et al. 2025). Herein, the hydrothermal pre-treatment step was followed by minimal mechanical processing consisting of a short (5 min) shredding process (**Figure 3a**). This procedure was applied to all selected plant tissues and another common food by-product, namely pomace from date processing (DP). For all samples, gels reminiscent of conventional cellulose nanofibers were obtained, where the presence of residuals resulted in colors in all gels except the watermelon mesocarp-based gels (**Figure 3b**).

**Figure 3.**
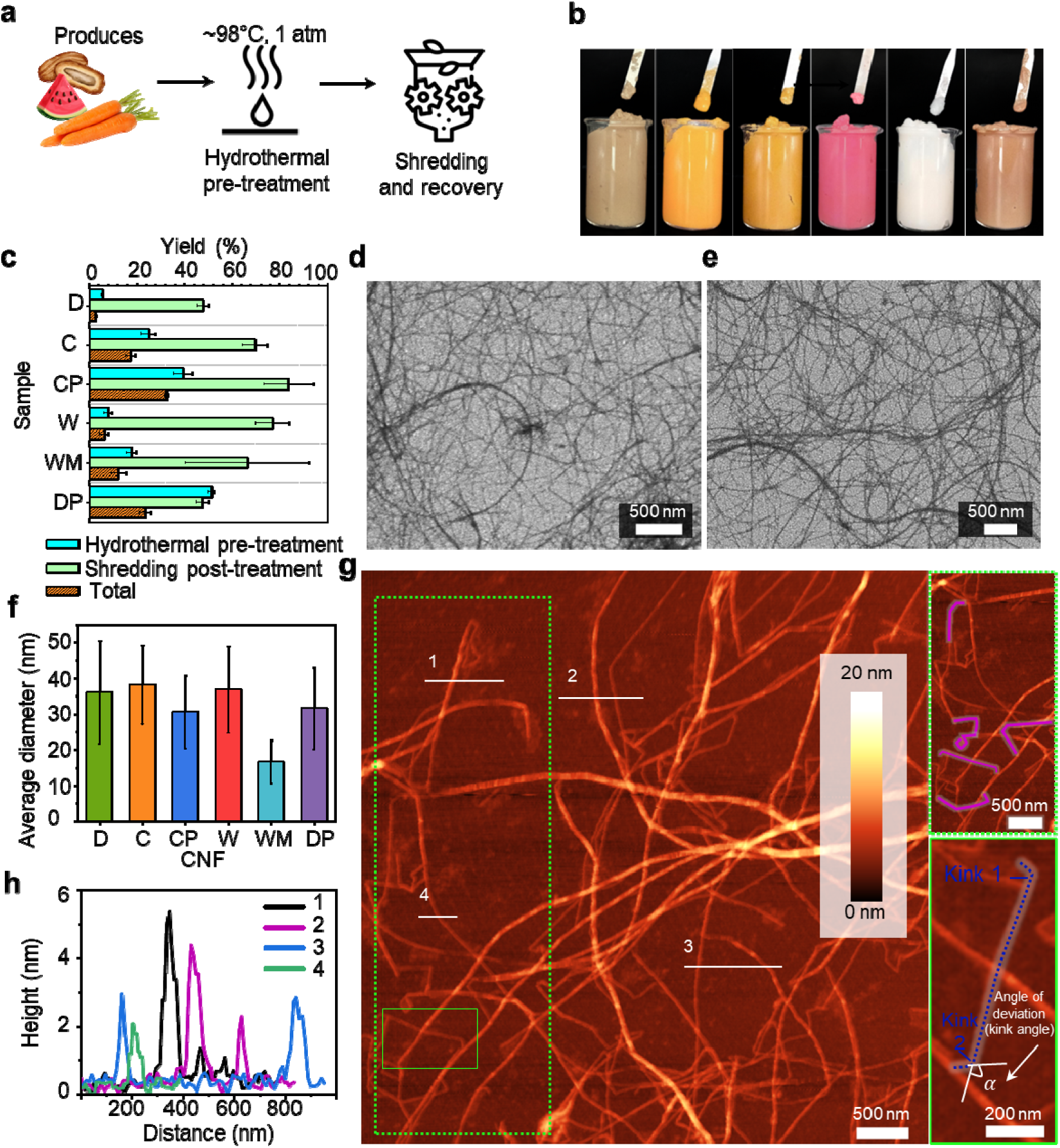
Nanofibrillation of non-structural plant tissues and resulting elementary fibrils containing CNFs. (a) Schematic illustration of the process of extraction and fibrillation of cellulose nanofibers. (b) Obtained CNF gels (D_CNF_, C_CNF_, CP_CNF_, W_CNF_, WM_CNF_, and DP_CNF_, from left to right, respectively). (c) Yield (%, dry weight basis) after hydrothermal pre-treatment, shredding posttreatment, and total (cumulative) process of CNF isolation; (d,e) SEM images of the obtained (d) WM_CNF_, and (e) D_CNF_. (f) Average diameter of CNFs as a function of plant tissue. (g) Atomic force microscopy (AFM) micrograph of WM_CNF_. Green contoured areas showcase kinks along fibrils’ axes (purple and blue lines). White horizontal lines are associated with (h) that shows the corresponding height profiles obtained for six fibrils, reaching elementary fibrils diameters.

Yields (dry weight basis) after “hydrothermal pre-treatment” followed by shredding and CNF recovery “shredding post-treatment” were obtained for each sample, along with the overall extraction yield, “total” (**Figure 3c**). The highest hydrothermal pre-treatment yields were observed for date pomace (DP_Hydro_) and carrot periderm (CP_Hydro_), 51%, and 39%, respectively, while the lowest yields were W_Hydro_ and D_Hydro_ at 5.5% and 7.9%, respectively. All samples showed relatively high yields following the subsequent shredding and gel recovery step. The highest total yield was measured for CP_CNF_ at around 32%, followed by DP_CNF_ at 23%.

SEM images of the obtained CNF from date and watermelon mesocarp (**Figures 3d–e**) confirmed that the samples were nanofibrillated, without evident fractions that did not undergo sub-cellular sizes fractionation. Average diameters of the extracted cellulose nanofibers (bearing the additional ‘CNF’ subscript) (**Figure 3f**) were conducted based on the obtained SEM images. All CNFs, consisting of either elementary fibrils or bundles (Mattos et al. 2019), exhibited an average apparent diameter lower than 40 nm. WP_CNF_ yielded the finest fibers, with the lowest average diameter of 16.6 ± 6 nm observed (**Figure 3f**). D_CNF_ and W_CNF_ nanofibrils were larger, with average diameters of 36 ± 14.4 nm and 36.7 ± 12.1 nm, respectively (**Figure 3f**).

The nanoscale morphologies of CNF isolated from WM were further characterized using AFM (**Figure 3g**). CNFs derived from watermelon peels exhibited a high degree of fibrillation, where the height profiles showed diameters between 2.1 and 5.4 nm (**Figure 3h**), consistent with the presence of elementary nanofibrils as observed in other fibrillation methods (Beaumont et al. 2021). The presence of some fibers smaller (i.e. 2.1 nm vs 3.5 nm) than those previously reported in wood (Beaumont et al. 2021) was likely associated with the fact that cellulose may aggregate in a distinct manner in watermelon as a result of a high pectin content in the extracellular matrix. Though some reports suggest that such sizes can also be observed in wood CNFs, particularly in high pressure homogenization setups (Wu et al. 2021). The amount of residuals apparent around the fibers was surprisingly small compared to wood, where up to 20% residuals are associated with hemicelluloses (Xiang et al. 2019), which typically form nanoparticles within the wood-CNF gels (Mattos et al. 2019). Only a few polymeric residues could be observed surrounding some of the fibers sections. More remarkably, the elementary fibers, typically containing alternating crystalline and amorphous domains (Tardy et al. 2021), showed an arrangement into sharply angled segments, so called kinks (**Figure 3g**, *right)* as previously reported for elementary fibrils isolated by chemical functionalization (Usov et al. 2015; Tardy et al. 2021; Zhen et al. 2024). These angular deviations can be quantified by measuring the kink angle (α), defined as the interior angle formed between two linear segments of the fibril at the bending point (**Figure 3g**, *bottom right*) (Zhen et al. 2024), and can originate from both structural and mechanical factors. Early studies proposed that kinks occur at the amorphous domains of cellulose, where the fibril chains are less ordered and thus more susceptible to local bending or slippage under stress (Usov et al. 2015; Nyström et al. 2018; Zhen et al. 2024). However, further investigations suggested that mechanical treatments such as homogenization and sonication can also induce kink formation even within crystalline segments (Usov et al. 2015), producing sharp angular distortions due to transient mechanical strain or interlayer slippage between crystal planes (Zhen et al. 2024). According to Zen et al (Zhen et al. 2024), higher-crystallinity cellulose tends to form kinks rather than undergo smooth bending or fracture, since its rigid crystalline regions resist longitudinal splitting and thus concentrate strain at limited sites, producing distinct angular defects. Together with the apparent absence of residuals, this convincingly demonstrates that the highest grade of nanocellulose can be isolated herein without affecting the paracrystallinity of cellulose, as the process is mild, nor does it alter its surface functional groups as is commonly done to obtain elementary fibrils. These observations further support that in plant tissues where cellulose serves a non-load-bearing role, the nanofibrillar network is sufficiently open to enable hydrothermal pre-treatment combined with shredding as a nanofibrillation method.

A more comprehensive comparison across all six samples is provided in **Figure 4**, which includes detailed SEM micrographs and corresponding diameter distribution histograms. While all CNFs consisted of either elementary fibrils or small bundles, differences in uniformity and dispersion were evident. WM_CNF_ exhibited a narrower diameter distribution, indicating more homogeneous fibrillation (**Figure 4l**). DP_CNF_ and average diameter comparable to CP_CNF_, however DP_CNF_ displayed greater variability in diameter, consistent with its broader distribution curve (**Figure 4m**), and its SEM images revealed the presence of residual particles that were not fully removed during processing (**Figure 4f** *Circled*). D_CNF_ and W_CNF_ displayed relatively larger fibril sizes (**Figures 4h, 4k,** respectively), with average diameters of 36 ± 14.4 nm and 36.7 ± 12.1 nm, respectively, along with wider distributions (**Figures 4a, 4d,** respectively), suggesting less uniform fibrillation compared to WM_CNF_. C_CNF_(**Figure 4b**) showed intermediate behavior, with a moderately narrow distribution and fibril (**Figure 4i**).

**Figure 4.**
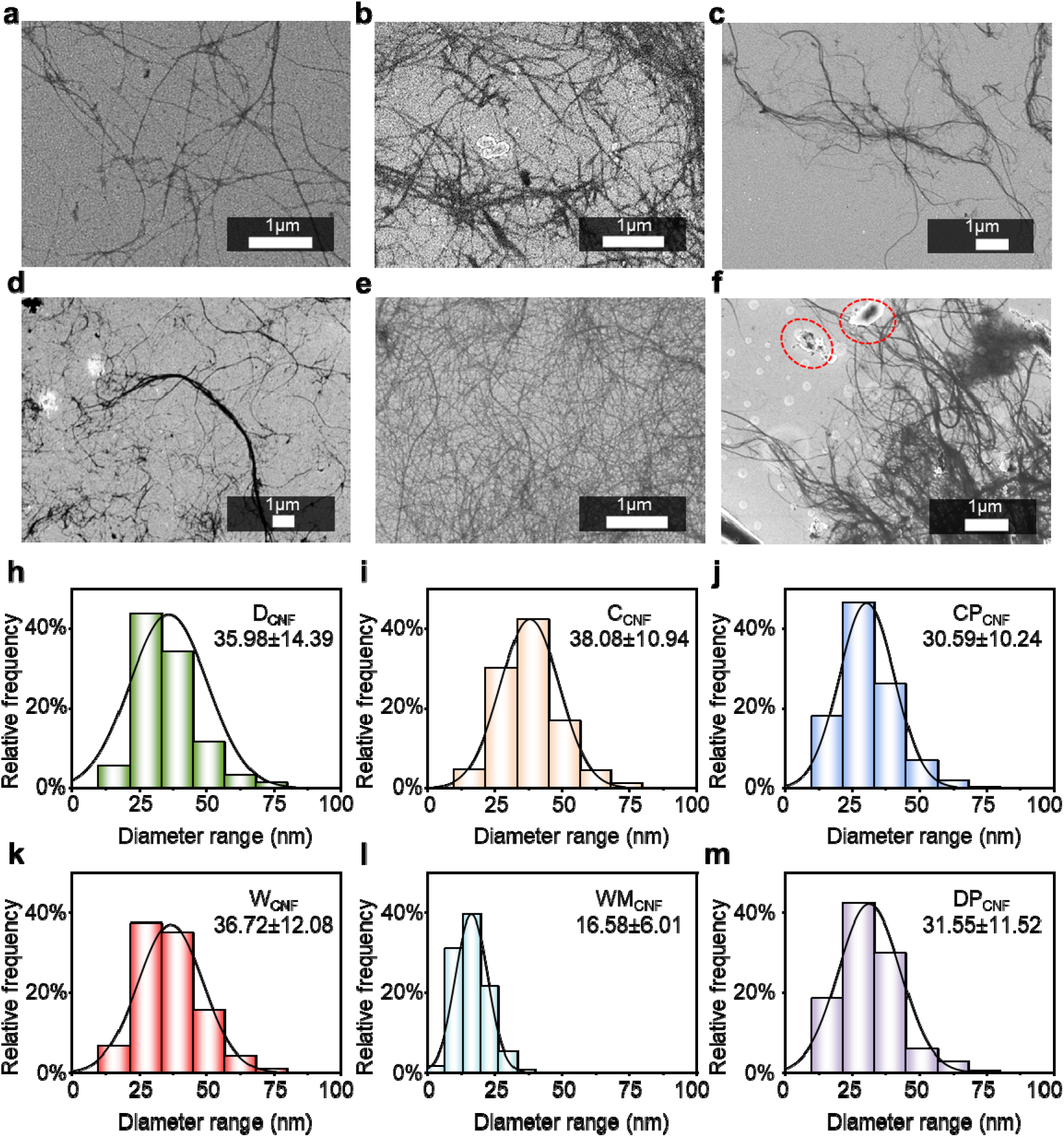
SEM images of the obtained (a) D_CNF_, (b) C_CNF_, (c) CP_CNF_, and (d) W_CNF_. (e) WM_CNF_, and (f) DP_CNF_; Size distribution histograms of (a) D_CNF_, (b) C_CNF_, (c) CP_CNF_, and (d) W_CNF_. (e) WM_CNF_, and (f) DP_CNF_, showing the diameter range of the nanofibers.

X-ray diffraction (XRD) analysis of the obtained CNFs from 6 different non-wood sources revealed three distinct peaks at 2θ ≈ 16.3°, 22.3°, and 34.8°, corresponding to the (110), (200) and (004) crystalline planes, typical of plant cellulose I allomorph (**Figure 5a**) consistent with the reference microcrystalline cellulose pattern (**Figure S4**) (Shi et al. 2024). The apparent crystallinity index (CrI) of the CNFs, calculated using Segal’s method, showed that WM_CNF_ exhibited the highest crystallinity (∼62.8%), followed by C_CNF_ (52.6%), D_CNF_ (46.7%), W_CNF_ (44.0%), CP_CNF_ (38.7%), and DP_CNF_ (36.1%). Removal of crystal forming non-cellulosics was apparent in all samples, when compared to untreated samples (bearing an additional 0 subscript) (**Figure 5a**) – particularly considering the reduction of the broad amorphous hump between 2θ ≈ 15° and 22.5°. FT-IR spectra of the processed samples displayed characteristic cellulose bands, including O–H stretching (3362 cm^−^¹), C–H stretching (2922 cm^−^¹), and C–O–C/C–O vibrations (1000-1200 cm^−1^) (**Figure 5b-g and S5**) (D’Orsi et al. 2023; Rasheed et al. 2024; Sharma et al. 2025; Tsaousis et al. 2025). The FT-IR spectrum of the processed samples (**Figure 5b-g**) also shows the disappearance of lignin-associated peaks at 1580, 1490, and 1450 cm□¹ (Zhang et al. 2023; Kostryukov et al. 2023), as well as the aromatic C–H out-of-plane bending at 830-870 cm□¹ (Derkacheva et al. 2021), along with the loss of the hemicellulose-related C=O stretching at 1730-1740 cm□¹ (**Figure 5e and 5g**) (Pancholi et al. 2023), while retaining the cellulose bands, confirming the effective removal of lignin and hemicellulose and the preservation of the cellulose structure.

**Figure 5.**
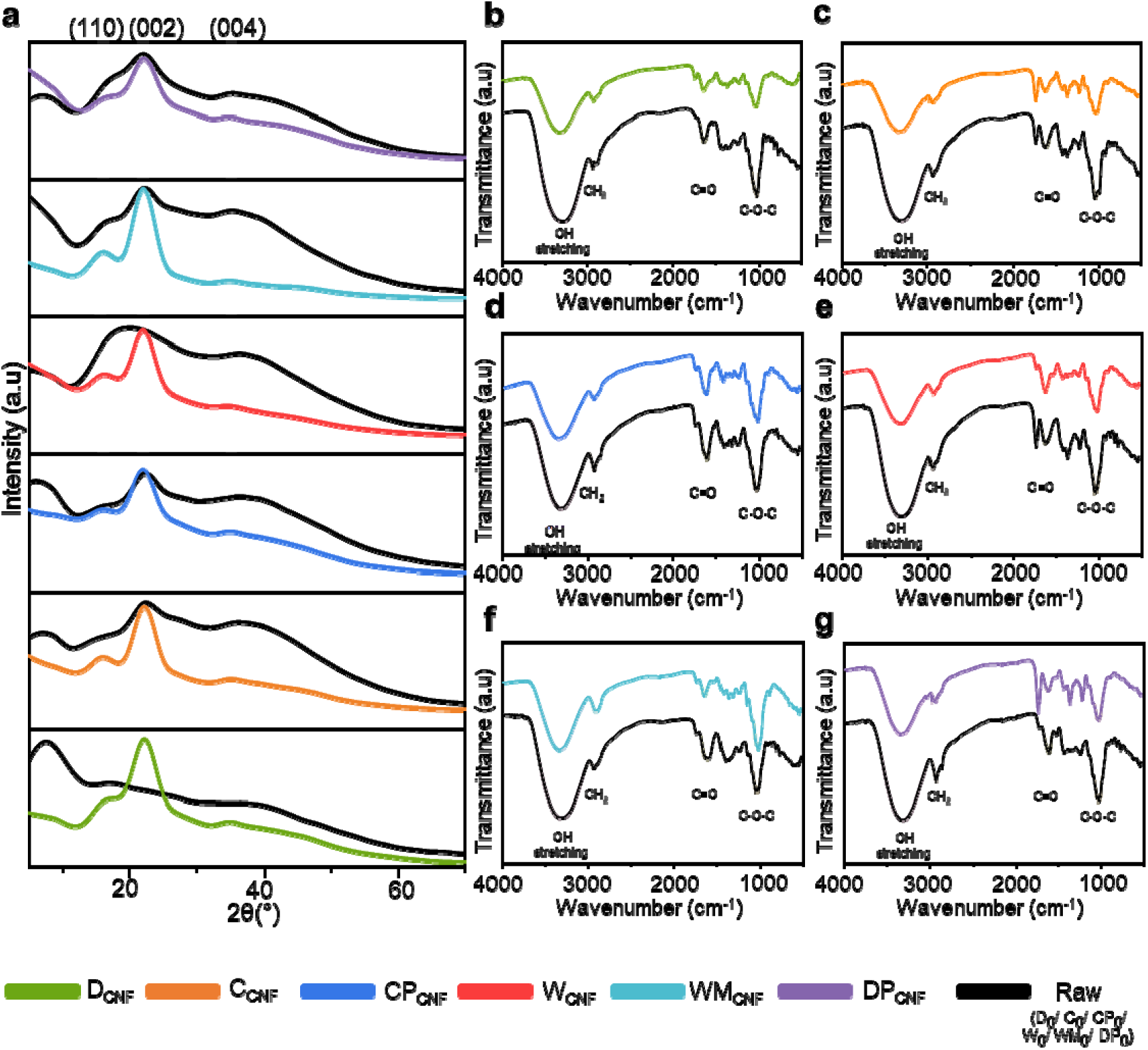
(a) X-ray Diffraction (XRD) spectra of raw (black) and processed biomass (colored). CNFs Fourier Transform InfraRed (FT-IR) spectra of (b) D_CNF_, D_0_; (c) C_CNF_, C_0_; (d) CP_CNF_,CP_0_; (e) W_CNF_,W_0_; (f) WM_CNF_, WM_0_; and (g) DP_CNF_, DP_0_.

To investigate the fibers’ thermal decomposition and mass loss profiles, TGA analysis was performed. The TGA curves of the obtained samples (**Figure 6a**) display the thermal degradation behavior of the cellulosic materials, showing typical three-stage weight loss pattern. The initial minor loss below 110°C corresponds to moisture evaporation, the major degradation between around 250°C and 380°C corresponds to cellulose depolymerization and glycosidic bond cleavage, and the residual mass above 400°C represents char formation and degradation of the remaining inorganic residues (González Martínez et al. 2022; D’Acierno et al. 2023).

**Figure 6.**
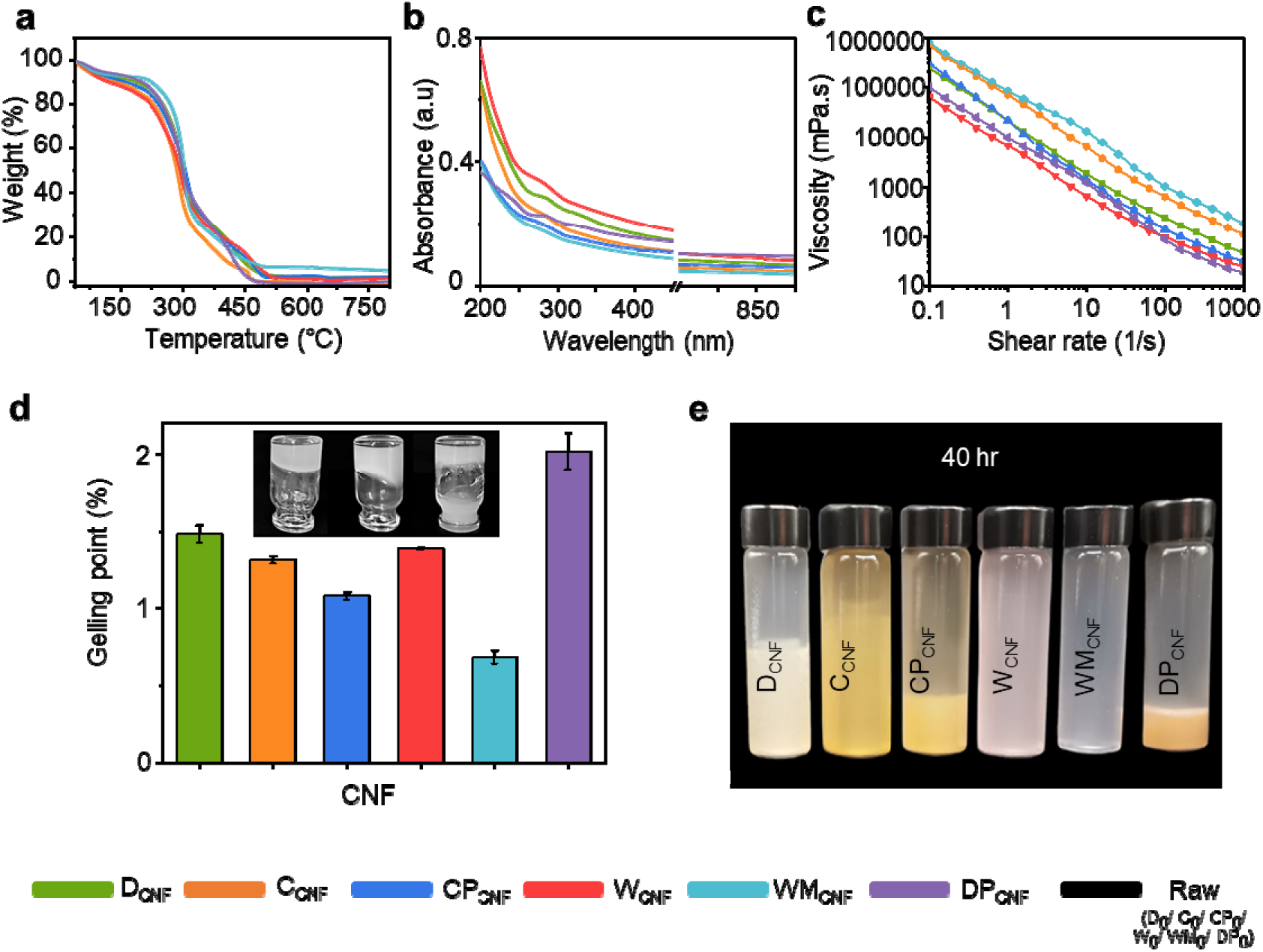
(a) Thermogravimetric assays (TGA). (b) The ultraviolet-visible (UV–vis) transmittance spectra, and (c) Rheological properties of the obtained CNF. (d) Gelling point concentrations of the obtained gels (inset image: gelation test jar of WM_CNF_. (e) Sedimentation near equilibrium heights at 0.25% initial concentration CNF (jars of D_CNF_, C_CNF_, CP_CNF_, W_CNF_, WM_CNF_, and DP_CNF_, from left to right, respectively after ∼40 hours test).

UV-Vis spectroscopic analysis (**Figure 6b**) indicated that CP_CNF_ and WM_CNF_ did not show specific adsorption peak between 200 and 900 nm, wherein only a slight shoulder could be observed at 280 nm. A clearer adsorption band was more clearly observed for DP_CNF_, D_CNF_, and W_CNF_, which is more likely associated with phenolic residues than with proteins, given the original nature of the tissues. Absorbance intensity at 900 nm can be attributed to scattering, which is related to the opacity of the dispersion; as expected, WM_CNF_ had the weakest absorbance. For all samples, it is likely that a set of residuals was convoluted, leading to the absence of clear adsorption bands.

UV spectra of the post-centrifugation supernatant were also analyzed (**Figure S6a**). At around 200-220 nm, all samples showed a dominant peak, with the highest intensity observed for the date and date pomace supernatants. Another pronounced peak at around 280 nm was also evident in these same samples (D_CNF_ and DP_CNF_). These peaks were attributed to π→π* transition in aromatic compounds such as residual phenols, tannins or small lignin-derived fragments, that contain chromophoric groups capable of absorbing in this range (Hynynen et al. 2021). At around 460 nm, a distinct peak was observed mainly in the UV spectra of carrot and carrot peels supernatants, but not in those of the other samples. This peak most likely corresponds to carotenoids (β-carotene), a characteristic pigment in carrots (Sricharoen et al. 2016). These results confirm that non-cellulosic compounds and other water-soluble extractives, commonly present in fruit wastes, were further removed during the centrifugation step, as shown in the SEM image of the W_CNF_ supernatant, with fine fibers also observed (**Figure S6b**).

To study the flow behavior of the gels, shear stress was measured as a function of shear rate, and the viscosity vs. shear rate curves were subsequently constructed (**Figure 6c**). All samples exhibited shear-thinning behavior, as expected for CNF suspensions (Li et al. 2021a). At low shear rates, WM_CNF_ showed the highest viscosity, while W_CNF_ showed the lowest, suggesting that WM_CNF_ possesses a more strongly entangled CNF network compared to the others. Up to a shear rate of around 1 s^−1^, all samples showed comparable slopes, indicating similar initial flow behavior. However, at high shear rates, DP_CNF_ demonstrated a steeper profile, reflecting greater sensitivity to shear and ultimately reaching a lower final viscosity than W_CNF_ and the other samples.

The microstructure of viscoelastic materials determines the form of the frequency sweep curves, which were studied herein at a strain of 0.01% (**Figure S7a**) (Li et al. 2021a), as determined from the amplitude sweep test (**Figure S7b**). Over the entire strain range, both WM_CNF_ and C_CNF_ exhibited storage modulus (G′) values higher than the loss modulus (G″), indicating an elastic dominant gel with a stable structure. Moreover, the G′ curve of both samples was nearly flat and frequency independent, suggesting stable gel-like materials with strong fiber-fiber interactions(You et al. 2025). The highest stability was observed for WM_CNF_, with dynamic moduli reaching approximately 3700 Pa and 950 Pa for G′ and G″, respectively. An important factor influencing the storage modulus of cellulose nanofiber gels is the aspect ratio of the fibers. Materials with a high aspect ratio tend to form extensive entanglements, which restrict fiber mobility under flow and lead to the formation of a stable, interconnected network, resulting in a higher storage modulus (Czaikoski et al. 2020). This further suggested that WM_CNF_ and C_CNF_ based gels have higher aspect ratio fibers. D_CNF_, W_CNF_, CP_CNF_, and DP_CNF_ showed a higher G′ than G″; however, at higher frequencies, a crossover was observed, reflecting disruption of the gel network under faster oscillations and indicating the presence of a sol-gel transition or a weak gel structure that breaks down at higher frequencies. Across the entire temperature range (20–80□), all gels exhibited storage moduli significantly higher than their corresponding loss moduli, confirming that the systems maintain a predominantly elastic, gel-like character (**Figure S7c**). Fluctuations, mainly in G″, were observed primarily in D and DP based suspensions. Such instability may arise from residual components or impurities that interfere with fibril-fibril interactions, thereby destabilizing the network and leading to more sensitive viscous response to temperature variations (Nechyporchuk et al. 2016). This suggests the presence of aromatic or phenolic compounds, reflecting their higher gelling thresholds (2.0% and 1.48%, respectively) (**Figure 6d**).

WM_CNF_ exhibited the lowest gelling point concentration, 0.68%, as determined by tube inversion assays (**Figure 6d**). This low gelling point could be attributed to a highly fibrillated CNF structure characterized by a small fiber diameter, as also suggested by SEM and AFM micrographs analysis (**Figure 3 e and g**). An increased interaction with water was unlocked with the higher surface area (Dali et al. 2025) of the WM_CNF_ and their high aspect ratio led to gelation at low concentration (Arcari et al. 2020), below what is typically observed for wood nanocelluloses. DP_CNF_ showed the highest gel concentration at ca. 2.0% followed by D_CNF_ at ca. 1.48%. As previously suggested by their UV results, aromatic or phenolic compounds might be present, reflecting their higher gelling thresholds. A similar trend was also observed in the gelation behaviour of W_CNF_, which exhibited a gelling point close to that of date at 1.39%. Differences in gelation concentrations, which cannot be directly associated with measured diameters, were likely related to floc size and the individualization of nanofibers (Raj et al. 2016). Residuals, such as phenolic compounds, may also play a role as they can alter fiber-fiber interactions.

To examine the dispersion stability of the six gels, stability assays were conducted by sedimentation over prolonged periods (**Figures 6e and S7d**). Within the first 7 minutes, a rapid 80% sedimentation was observed in DP nanofibers, forming a sedimented gel layer at the bottom and leaving an almost clear supernatant above. This was followed by an equilibrium phase, during which no further change in height occurred for the remainder of the test. CP nanofibers followed a similar pattern, with 70% of the sedimentation occurring in two stages: an initial rapid sedimentation of 60% within the first 15 minutes, followed by 10% that occurred at a lower rate. Afterward, the sedimentation height stabilized, leaving a visibly turbid supernatant that indicated partial fiber suspension. The CNFs derived from W and its mesocarp (WM) demonstrated the highest dispersion stability, as they did not sediment throughout the 24-hour test. The improved colloidal stability can be attributed to the presence of kinks, as indicated in the previous AFM images, that create steric hindrance preventing neighbouring fibrils to come into close contact and forming heavier bundles (Usov et al. 2015). C exhibited gradual sedimentation over multiple stages. Less than 10% occurred within the first 5 minutes, and the test ended with less than 25% of partial sedimentation. Among the samples, date-derived CNF showed moderate stability, with the sedimentation front remaining at ca. 50% (**Figure S7d**). Together, the combined FTIR, XRD, AFM, and SEM observations consistently indicate a significant enrichment in cellulose content and a pronounced reduction in non-cellulosic phases.

To explore the potential of the extracted cellulose nanofibers for material applications, the prepared gels were tested for their film-forming ability. Film morphology (**Figure 7a**) enabled relative performance in the dried state to be evaluated. The appearance of the films was smooth, and the colors reflected those of the gels. Macro-aggregates were visible on all films, suggesting that a fraction of large fibers remained. WM_CNF_ and DP_CNF_ films were shown to form nanolayered stacks of nanocellulose, typical of so-called nanopaper (**Figures 7b, S8-S12**) but only WM_CNF_ presented a smooth surface, as associated with large unfragmented residuals (**Figures S8 and S9**). Stress-strain behavior (**Figure 7c**) revealed significant variation in the mechanical properties depending on the feedstock. Of note, all films were prepared, processed, and tested under identical conditions.

**Figure 7.**
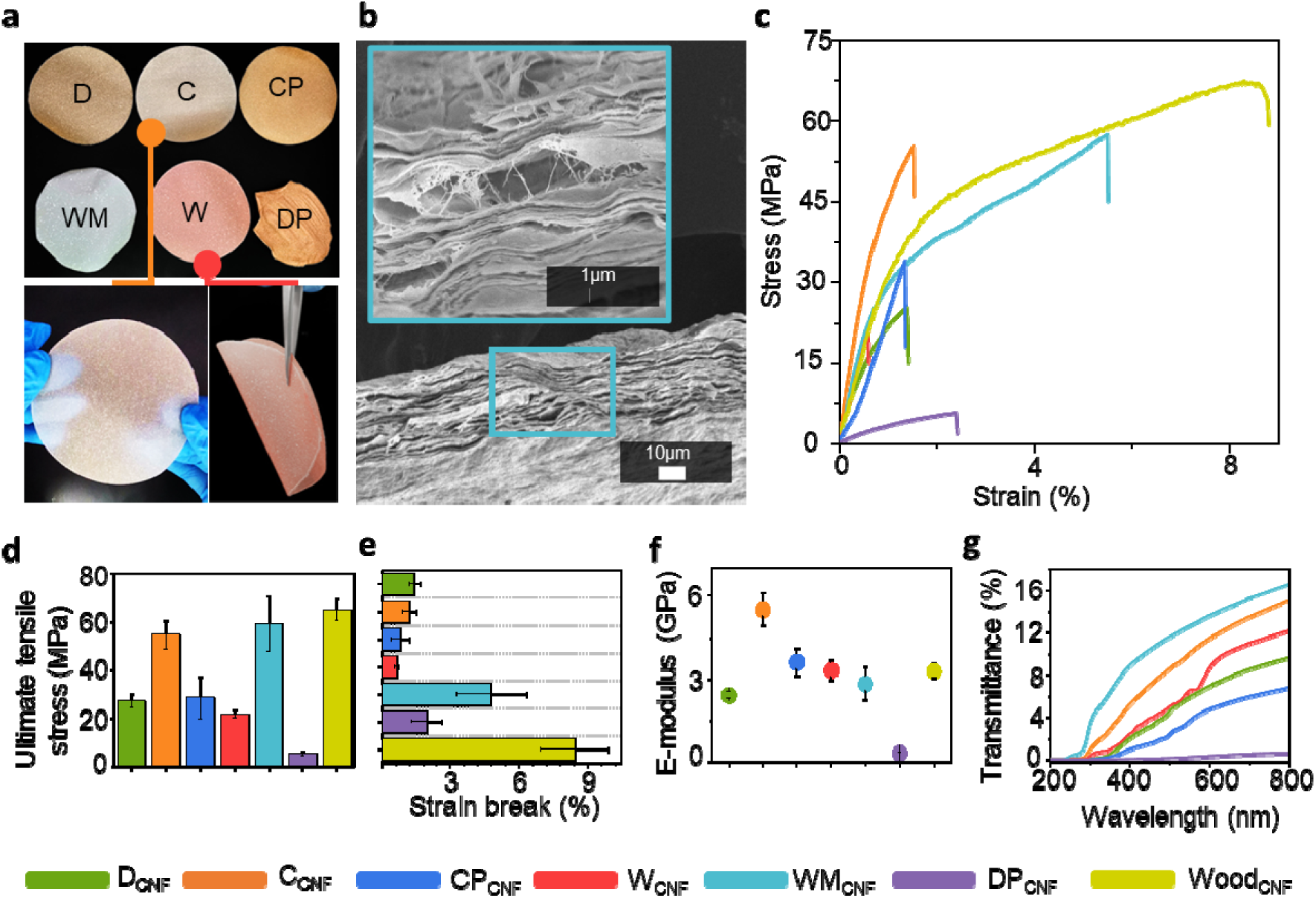
Benchmarked performance of nanopaper obtained from hydrothermally obtained CNFs. (a) Prepared films (nanopaper) obtained from the respective CNFs (top) with the bottom panels showcasing transparency and flexibility of the obtained films (C_CNF_ (*left)*, W_CNF_ (*right*) film). (b) Cross-sectional SEM of nanopaper obtained from WM_CNF_. (c) Representative tensile stress-strain response of films, (d) Ultimate tensile strength, (e) Strain at failure, (f) Elastic modulus of the CNF films obtained from the non-wood biomass and wood. (g) UV-vis transmittance spectra of the films.

While most samples exhibited primarily elastic deformation until failure at strains below 1.5%, typical of nanocellulosic materials, the WM_CNF_ film showed plastic deformation extending beyond 5% and up to 16% in some samples. This ductility, which exceeds values typically reported for nanocellulose films, could be attributed to the presence of kinked fibrils previously observed in AFM images (**Figure 3g and i**), which act as flexible joints within the network. As illustrated by Zhen et al. (Zhen et al. 2024), during loading, kinks gradually straighten, dissipating stress and thereby allowing higher elongation before failure. Highly curled nanocellulose with abundant kinks also forms more interlocked networks that enhance toughness and strain at break (Zhu et al. 2015; Walther et al. 2020; Zhen et al. 2024). Beyond the presence of kinks, the high strain observed can also be attributed to ambient humidity(Walther et al. 2020) and the structure of cellulose fibers (Zhu et al. 2015). Higher relative humidity results in moisture being absorbed by the CNF films, allowing fibrils to slide more easily relative to each other and thus increasing the overall strain while moderately lowering stiffness (Walther et al. 2020). At the molecular level, cellulose nanopaper under tension experiences a cascade of hydrogen bond breaking and reforming event where this reversible bond dynamics enables progressive stress relaxation and plastic-like elongation without rupture (Zhu et al. 2015).

Ultimate tensile strength (UTS) (**Figure 7d**) was highest for WM_CNF_ and C_CNF_ films (>50 MPa), followed by CP_CNF_, D_CNF,_ and W_CNF_ (∼22-28 MPa), while DP_CNF_ films exhibited the lowest strength (5 MPa). Despite its low strength, DP_CNF_ films appeared to display moderate strain values (2%) (**Figure 7e**), which is likely to be associated with macro-scale wrinkles in the sheets (**Figure S10**). Once these wrinkles were straightened under load (**Figure S12a**), fracture occurred rapidly, reflecting weak intrinsic bonding within the DP nanofibril network (**Figure S12b**).

While the tensile strength generally correlated with crystallinity, WM_CNF_, being the most crystalline, maintained moderate stiffness (E ≈ 3300 MPa), lower than C_CNF_ and CP_CNF_ (E ≈ 5500 MPa and 3600 MPa, respectively) (**Figure 7f**). However, WM_CNF_ and C_CNF_ films exhibited both relatively high strength and strain, resembling the behavior reported by Zhu et al. (Zhu et al. 2015), where the smaller the fibers, the stronger and the tougher. This was explained by fewer defects and more uniform fibrils, leading to a more even stress distribution and reduced crack initiation. Smaller diameter fibers provide a larger surface area, thus a higher density of hydrogen bonds, resulting in improved strength and toughness (Zhu et al. 2015).

This pointed to an interplay of multiple microscale phenomena affecting the overall mechanical behavior of the films. For instance, SEM and gelation analyses showed that WM_CNF_ CNFs were highly fibrillated, with a low gelation concentration, enabling strong inter-fibril entanglement. This was also consistent with its rheological profile, in which its high storage modulus indicates a stable, elastic-dominant gel network with the capacity for efficient stress dissipation. Conversely, the poor mechanics of DP_CNF_ correlate with its low crystallinity (Ottesen et al. 2019), partial fibrillation observed in SEM, and high levels of residual phenolics that interfere with the inter-fibrillar hydrogen bonding resulting in lower structural integrity of films. To benchmark the prepared hydrogels and mitigate the impact of preparation or measurement methods that typically lead to a wide range of values for similar samples, a wood-derived CNF film, using previously reported precursors (Österberg et al. 2013; Toivonen et al. 2015; Mattos et al. 2019), was also prepared (**Figure S13**) and tested (**Figure 7c**). It exhibited an ultimate tensile strength of approximately 65 MPa (**Figure 7d**), a strain at break of about 8% (**Figure 7e**), and a modulus of approximately 3300 MPa (**Figure 7f**). These reference values indicated that the non-wood CNF films developed here, particularly those from WM and C sources, achieved comparable mechanical properties, even though they were obtained through significantly milder processes. These results collectively highlighted the interplay between fibril morphology (kinks, and size), hydrogen-bond dynamics, and moisture sensitivity, in governing the mechanical behavior of nanocellulose films.

A key performance property across various applications of nanocellulose-based films is their interaction with light, as quantified by UV-vis transmittance measurements (**Figure 7g**). All samples showed a monotonic increase in T (%) with wavelength, with differences in magnitude and slope among the films. In the visible region (400–800 nm), both WM_CNF_ and C_CNF_ films exhibited the highest transmittance (T%) exceeding 14%. This indicates their high optical clarity (**Figure 7a** *bottom left*) with superior nanofiber dispersion. However, DP_CNF_ films remained nearly opaque, with a transmittance of less than 2%. This significant light absorption was likely associated with poor dispersion, high surface roughness or degree of wrinkles (**Figure S10**), and the presence of residual contents or other UV-absorbing impurities. In the UV region (200–400 nm), DP_CNF_ and CP_CNF_ exhibited relatively low transmittance (<1%), indicating enhanced UV-blocking performance for potential applications in coatings and packaging.

The results obtained in this work underscore the relevance of these findings within real food systems. For example, cellulose nanofibers were observed in commercially available carrot purée (**Figure 8**), a product commonly used in baby food, demonstrating that such nanostructures are not limited to laboratory-processed materials but are already present in widely consumed food products.

**Figure 8.**
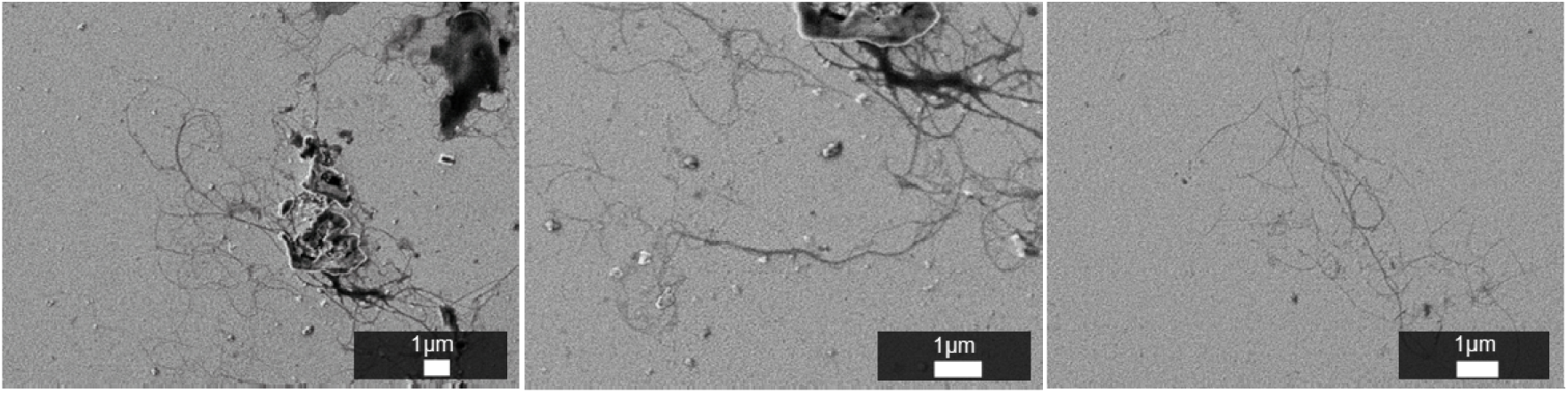
SEM images of CNF found in commercial carrot puree.

## CONCLUSION

The findings put forward in the manuscript open new directions for leveraging biological design in sustainable materials manufacturing. Given adequate plant ultrastructure, a mild hydrothermal treatment, combined with food-grade mechanical processing (i.e. shredding), enables the isolation of high-grade CNFs. The method applied to three plant tissues resulted in high-yields and high-grade CNFs. We demonstrate that successful nanofibrillation is largely associated with the tissues’ composition and, more importantly, their inherent ultrastructure. This approach as a high potential to be generalized to a wider range of tissues following the guidelines provided herein, i.e. as associated with low lignin and high water contents and importantly to the load-bearing functions of the tissues’ cells. The hazards-free CNFs are inherently natural nanomaterials, overcoming key legislative challenges associated with their integration across various sectors, including the food supply chain and environmental applications. The impact of the findings is expected to substantially change the isolation of nanocellulosics and their integration across the food chain, as the findings imply that nanocelluloses are already an integral part of food products.

Among the obtained samples, WM_CNF_ exhibited the smallest average diameter (<17 nm, with elementary sizes of 4 nm or below), and the lowest gelation concentration (∼0.68%), outperforming the other five tissues evaluated. Given the use of water only as a solvent and the low energy input of the applied mechanical step, the production cost of such CNFs is also expected to decrease substantially. Thus, the results of this study open a wide range of opportunities for (i) enhancing and advancing our understanding of nanocellulose across diverse plant tissues, (ii) creating new avenues for the scientific community to connect processes to each plant’s compositions and ultrastructure, (iii) applying low-cost and hazard-free process towards a better integration into the global circular bioeconomy of non-woods. Overall, this will engage a broad community of scientists to further explore this new approach to CNF isolation and its applications, which will overcome the limitations of costs and safety. Beyond the practical considerations above, this study opens the door to a more efficient global bioeconomy by facilitating the isolation of CNFs as high-performance precursors for forest-scarce regions (Tardy et al. 2023).

## EXPERIMENTAL SECTION

### Materials

Three types of cellulose feedstock were used in this study, carrots, watermelon, dates as well as date pomace. Carrots (Australian imported), Khalas dates, and readily available watermelon were purchased from a local market in Abu Dhabi, UAE. Date pomace was provided by Al Barakah Dates Factory located in Dubai, UAE. Deionized (DI) water (R: 15.0 MΩ cm) produced using a water purification system (Elix® - Merck Millipore) was used throughout the study. Tert-Butyl Alcohol (TBA, ACS reagent, ≥99.0%) was purchased from Sigma Aldrich. Polyvinylidene fluoride (PVDF) membrane filters, with a 0.2-micron pore size and 90 mm diameter, were purchased from Sterlitech.

### CNF extraction

CNFs were extracted from carrots, watermelon, dates, and date pomace by hydrothermal pre-treatment followed by shredding. Initially, thin slices of carrot peels (periderm) and the inner layers of flesh bypassing the core were separately collected using a scraper or peeler. An excess amount of DI water at room temperature was added to known mass of slices and heated to bring the mixture to boil. Since then, it was allowed to continue boiling, after which the used water was discarded and replaced with fresh DI water. The mixture was returned to the hot plate to continue heating, with low to moderate bubbling. This step was repeated as needed. Finally, the treated slices were strained from the water and allowed to cool to room temperature. A similar process was applied to watermelon flesh and white rind (mesocarp, located between the green rind and pink flesh), but with low bubbling intensity to accommodate their delicate structure. A similar procedure was followed for dates; however, during the final cycle, the dates were immersed in freshly boiled water and then left to soak overnight at room temperature without continuous heating. For date pomace, boiled DI water at a 1:10 ratio (DP:DI water, w/v) was mixed with the pomace, covered with aluminum foil, and placed on a hot plate to maintain 80□ with continuous, slow stirring. While still hot, the pomace was filtered through a piece of cloth supported by a mesh sieve, then gently pressed to remove excess water. The process was repeated a second time.

After hydrothermal pre-treatment, the treated products and pomace were shredded using a 500W Kenwood blender (BLP16.150WH) at the lowest power setting for 5 min only. DI water at a 2:1 ratio (DI water to washed slices) was added prior to shredding to ensure a uniform, smooth shredding process. The resulting gels were centrifuged once at 5000 rpm for 15 min to remove excess water. However, multiple centrifugation cycles were performed on the shredded date and date pomace, with the gels being separated from black tannin residues (**Figure S14**) until maximum possible separation was achieved while ensuring optimum gel recovery. The solid content (%) and yield of the process were later measured for the collected gels.

#### Nanopaper preparation

To prepare thin films, 40 mL of 0.4 wt% CNF gels were filtered through PVDF membrane filters using a vacuum filtration setup (**Figure S15**). The resulting films were then left to dry overnight at room temperature and subsequently detached from the membrane. To ensure a smooth surface and uniform thickness, the films were hot-pressed at 80□ for 2 min using a heat press (IMPRESOMATIC, 38×38 cm).

### Characterization

The morphology of the pre-treated biomass was analyzed using a scanning electron microscope (JEOL JSM-7610F FEG-SEM) after freeze-drying, preceded by a 10 min immersion in TBA (Inoué and Osatake 1988; Mattos et al. 2019). The morphology and size of the nanofibers were also analyzed using SEM. A 0.001-0.005 wt% droplet of nanofiber suspension was placed onto a gold-coated silicon wafer and left to dry at room temperature overnight. SEM images were then captured at an accelerating voltage of 5 kV and an emission current of around 60 µA. At similar conditions, the surface and cross-sectional morphology of the thin films were also analyzed using SEM before and after mechanical testing. The diameters of the CNFs were then measured from SEM images using ImageJ (NIH, USA). Diameter values from over 100 individual fibrils were collected across several micrographs to establish a size distribution histogram. Surface topography of the fibers was also analyzed using atomic force microscopy (CSI NanoObserver 2 AFM). A 0.001 wt% droplet of WM_CNF_ and D_CNF_ was placed on silicon wafers. A cantilever with a nominal resonance frequency of ∼300□kHz was employed. The images were further analyzed using Gwyddion 2.69 software (Czech Metrology Institute, Czech Republic) as reported in the SI. Fourier-transform infrared (Bruker Vertex 80v FT-IR) was used to evaluate the chemical composition and functional groups of the CNF before and after treatment. The samples were crushed after oven drying using a mortar and pestle, then pressed using a hand-operated pellet press. X-ray diffractometer (XRD PANalytical Empyrean) was used to evaluate the crystallinity of the samples. Cu-Kα radiation was used to obtain diffraction data over a 2θ range of 5° to 80° at 45 kV and 40 mA. UV-vis absorbance spectra of both the supernatants and gels collected after centrifugation were measured using a LAMBDA 1050 UV/Vis/NIR spectrometer over a wavelength range of 200-900 nm, with 1 nm intervals. Prior to testing, the gels and their corresponding supernatants were diluted to 0.01 wt%. UV-vis transmission spectra of the films were acquired under the same conditions. Freestanding films were mounted directly in the beam path, with air used as the reference. All measurements were performed at room temperature. Thermogravimetric Analysis (TGA) was performed using a high-temperature TGA system (STA 449 F3 Jupiter series, Netzsch, Germany). Measurements were conducted on ∼5 mg specimens under an air purge flow of 30 mL/min, employing a heating program from 40□ to 1000□ at a constant rate of 5□/min. Derivative thermogravimetric (DTG) curves were obtained for the fiber samples to assess decomposition stages.

Rheological measurements of the extracted cellulose nanofibers were performed using a rotational rheometer (Anton Paar H-PTD 220 FOR MCR) at a gel concentration of 2 wt%. Measurements were conducted at 20°C using a parallel plate geometry with a diameter of 25 mm and a fixed gap of 1 mm. To determine the viscous behavior of the samples, shear flow tests were conducted over a shear rate range of 0.1 to 1000 s^−1^. Viscosity as a function of shear rate was recorded at 16 logarithmically spaced points. Amplitude sweep tests were performed to determine the linear viscoelastic region (LVR) within the strain range of 0.01–100% at a constant angular frequency of 1 rad/s. The LVR limit was identified, and a conservative strain amplitude of 0.01% was selected. Subsequently, frequency sweep tests were carried out in the range of 0.1-100 rad/s, within the LVR (strain of 0.01%), using 16 logarithmically spaced points, to determine the storage modulus (G′) and loss modulus (G″). To assess the thermal stability and structural changes of the CNF suspensions, temperature ramp tests were conducted from 20 to 80°C under a fixed strain of 0.01% and an oscillation frequency of 1□Hz.

### Sedimentation Assays

Fiber suspensions (0.25 wt% in 5 mL of DI water) were sonicated for 5 minutes to ensure uniform dispersion and improve dispersion stability. The suspensions were then left to settle undisturbed in glass containers for a full day. In order to compare the sedimentation profiles of the various fiber suspensions, all created with similar initial volumes, photographic images were recorded throughout the settling process.

### Gelling Point Test

A known mass and solid percentage (%) of wet fiber gel was placed in a container (glass jar). Water was added dropwise to the sample to dilute the mixture, and the mixture was mixed using a vortex mixer to ensure uniform distribution. The jar was then inverted for 1 min to observe the behavior under the effect of gravity. This procedure was repeated until the wet fibers lost their integrity and began to flow or fall apart. The concentration of solids (%) in the gel at this stage is called the gel point.

### Mechanics

Tensile testing was performed on films cut into standardized dogbone-shaped specimens with a narrow section measuring 20 mm × 3 mm. The measurements were conducted in the dry state using a universal tensile tester (Instron 5966, USA) equipped with a 10 kN load cell. Typically, three to five specimens per sample were tested.

The solid content (%) of the gels was determined by measuring their mass before (m_W_) and after (m_d_) drying in the oven at 50□ and overnight to evaporate all the moisture present in the samples:

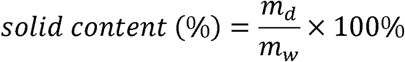

### Yield Calculations

Two yield values were obtained for each sample: (i) the yield of hydrothermal pre-treatment and (ii) the yield of shredding and centrifuging (mechanical processing and gel recovery). The overall yield of the CNF extraction process was then calculated as the product of the two yield values using the following equation.

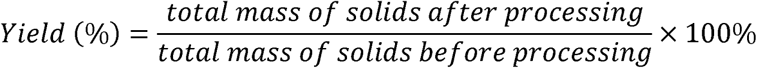

### Crystallinity Calculations

The crystallinity index (CrI (%)) was calculated using Segal’s equation (Segal et al. 1959):

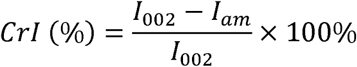

### Figure 1b Methodology

A multi-criteria framework was applied to compare the work done in this study with two routes based on energy efficiency, fibrillation performance, network mechanical performance, chemical safety, and cost efficiency. Both routes are wood-derived and start with bleached kraft pulp, followed by either (i) purely mechanical processing, or (ii) enzymatic pre-treatment using an endoglucanase (Novozym 476) followed by mechanical processing. The reference data for the present study corresponds to the WP_CNF_ sample. Energy efficiency was quantified as the total specific energy consumption (kWh·kg□¹) required to produce the equivalent of 1 kg dry CNF, including upstream kraft pulp production for wood-based routes and any subsequent pre-treatment and mechanical processing steps. Fibrillation performance was assessed using reported average nanofibril diameters, with smaller diameters indicating a higher degree of fibrillation. Network mechanical performance was evaluated using tensile strength values of pure CNF nanopapers reported in the literature, reflecting the integrity and load-bearing capacity of the fibrillar network. A scaling ratio was established based on the tensile strength values of bleached-kraft mechanically processed nanopapers from this work and those reported in the literature to avoid the effect of methodological differences among works. This ratio was used to scale the tensile strength of enzyme-CNF nanopapers. Chemical safety was assessed as the sum of products of reagent masses used per kg of CNF produced and the NFPA 704 health hazard ratings (0-4) of these reagents. The cost efficiency was determined by considering the costs associated with materials, reagents, and energy consumption. All parameters were transformed to a common dimensionless scale. The scale bounds are arbitrary and were chosen to graphically highlight the differences between the processes under comparison.

## Supporting Information Available

SEM images of hydrothermally treated carrot and watermelon tissues (Figures S1–S3); characterization of reference cellulose paper using XRD and FT-IR spectra (Figures S4–S5); analysis of the supernatant (UV-vis and SEM), (Figure S6); rheological behavior and sedimentation analysis (Figures S7); mechanical testing and cross-sectional morphology of CNF films (Figures S8–S13); additional nanofiber characterization and application-related images (Figures S14–S15).

## Acknowledgements

This research was funded by Khalifa University, KU-INT-RIG-2025-847400052, and the Food Security and Technology Center (FSTC), KU-INT-FSTC-2025-8474000576. The authors gratefully acknowledge the Research Council of Finland under the Genecellnano Flagship (No. 374296) for their support. The authors are thankful to Dr. Luiz Greca for his inputs and perspective associated with the manuscript. The authors also acknowledge the Center for Membranes and Advanced Water Technology (CMAT) at Khalifa University for access to facilities used in film preparation.

## Credit authorship contribution statement

**Malak AbuZaid**: Writing – review & editing, Writing – original draft, Methodology, Formal analysis, Data curation. **M-Haidar A. Dali**: Writing – review & editing, Methodology, Formal analysis. **Mohamed H. Salim**: Writing – review & editing, Methodology, Investigation, Formal analysis. **Vengatesan M. Rangaraj**: Writing – review & editing. **Marjo Yliperttula**: Writing – review & editing. **Fawzi Banat**: Writing – review & editing. **Blaise L. Tardy**: Writing – review & editing, Writing – original draft, Supervision, Methodology, Investigation, Funding acquisition, Formal analysis, Conceptualization.

## Notes

The authors declare that some of the findings are part of the patent application filed: PCT/IB2026/051222.

