## Supplementary information for "Isolation of Elementary Nanofibrils of Cellulose from Non-Structural Plant Cells: Hydrothermal Processing as a Generalizable Route"

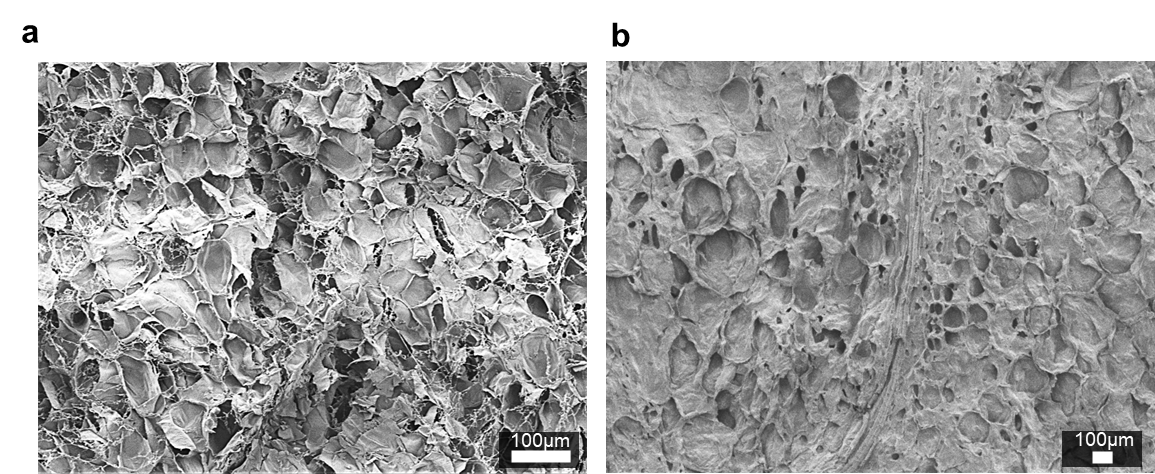

**Figure S1.** SEM of hydrothermally treated (a) carrot periderm (CP_Hydro_), and (b) watermelon mesocarp (WM_Hydro_).

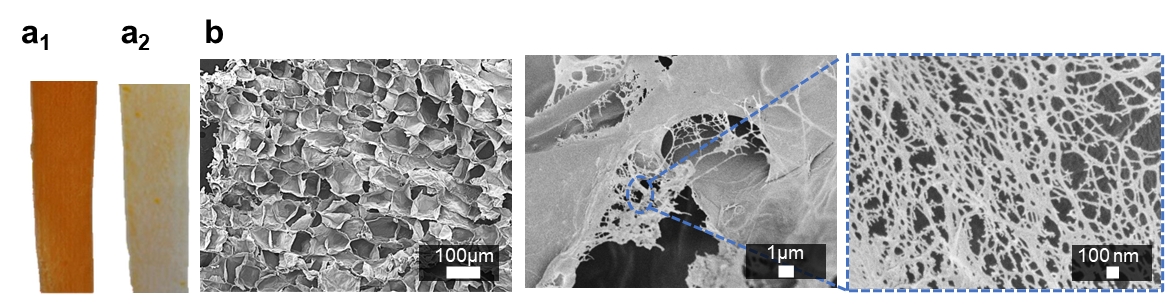

**Figure S2.** Carrot slice (a_1_) before (C_0_) and (a_2_) after hydrothermal pre-treatment (C_Hydro_); (b) SEM of C_Hydro_.

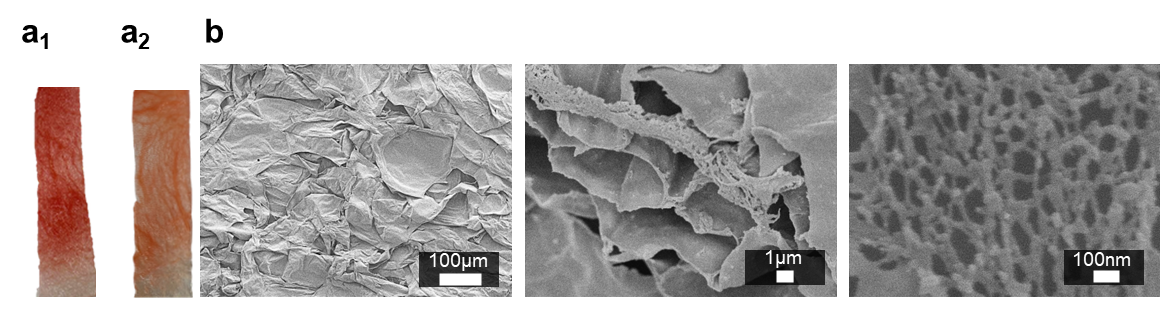

**Figure S3.** Watermelon slice (a_1_) before (W_0_) and (a_2_) after hydrothermal pre-treatment (W_Hydro_); (b) SEM of W_Hydro_.

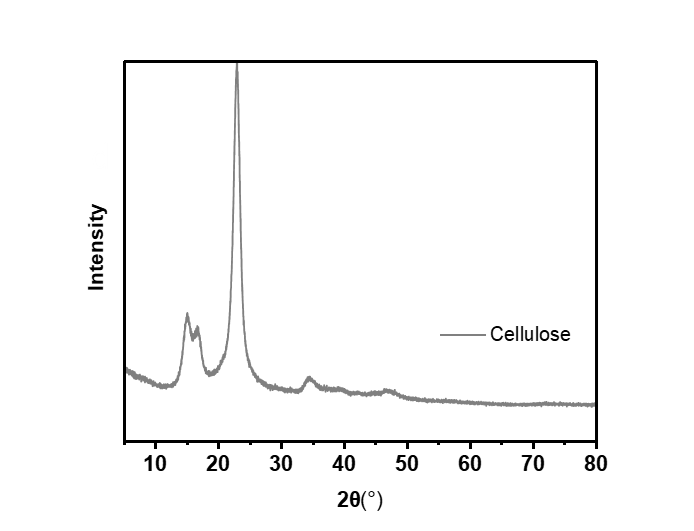

**Figure S4.** XRD spectrum of reference cellulose paper.

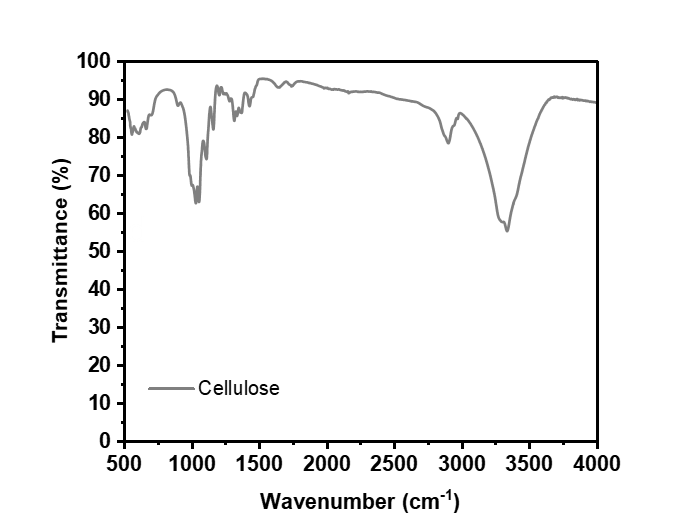

**Figure S5.** FT-IR spectrum of reference cellulose paper.

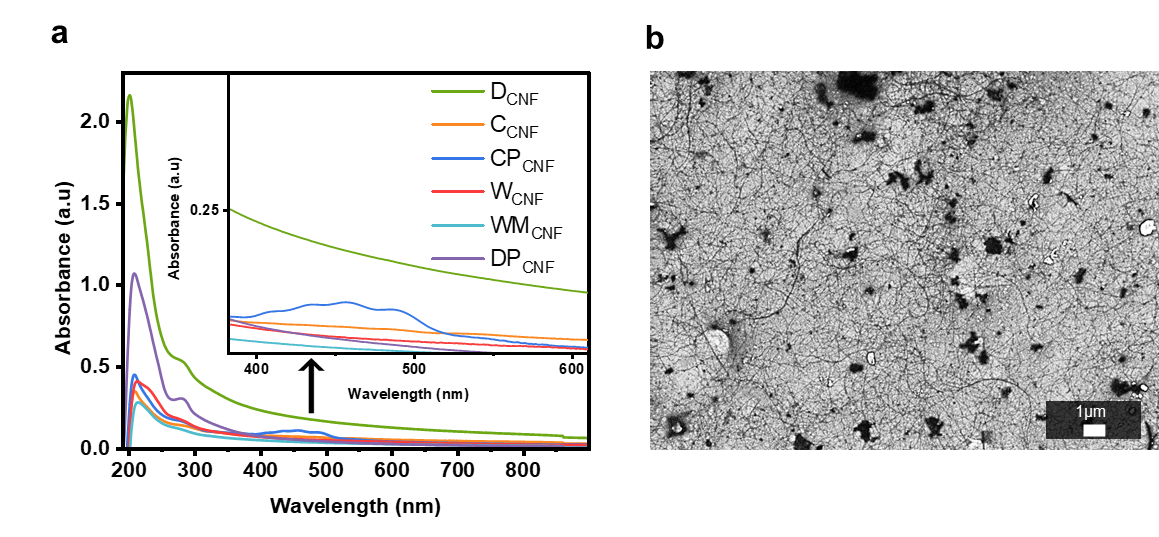

**Figure S6.** (a) UV-vis of supernatant of gels upon centrifugation; (b) SEM of supernatant from centrifugation of WM_CNF_ gel.

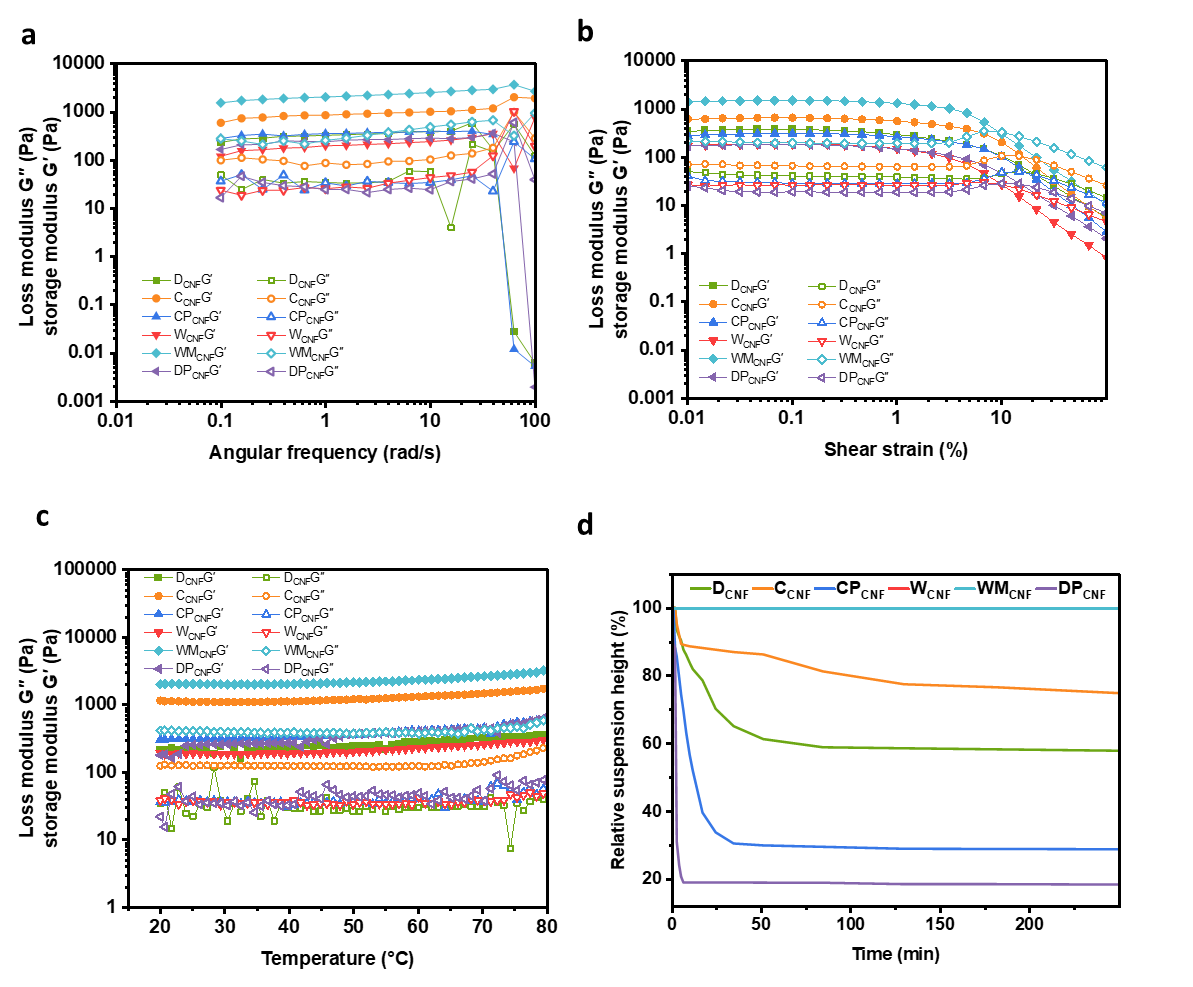

**Figure S7.** (a) Frequency sweep (b) Amplitude sweep and (c) Temperature ramp of the prepared CNF suspensions; (d) Relative sedimentation height (%) over time during sedimentation assays.

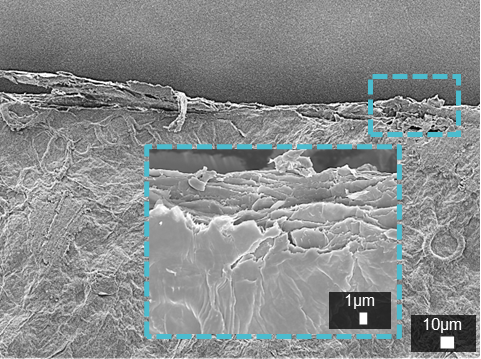

**Figure S8.** SEM image of the surface of WM_CNF_ film before mechanical testing.

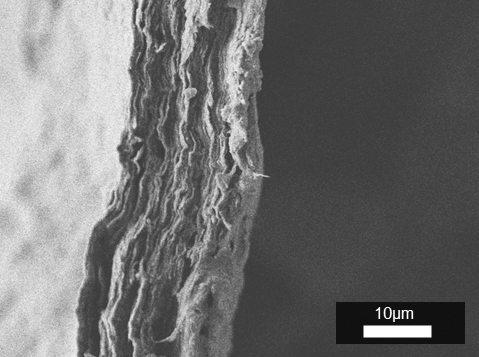

**Figure S9.** Cross-sectional SEM image of WM_CNF_ film after mechanical testing.

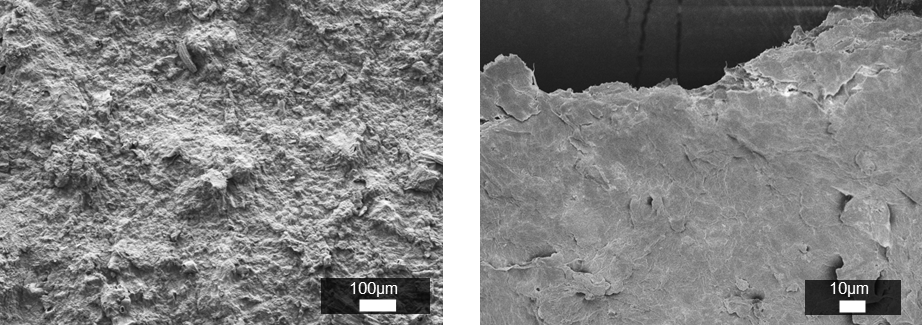

**Figure S10.** SEM images of the surface of DP_CNF_ film before mechanical testing.

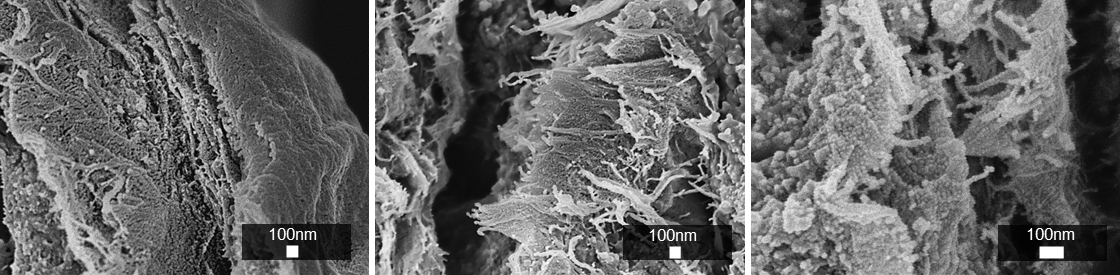

**Figure S11.** SEM images of the cross-section of DP_CNF_ film before mechanical testing.

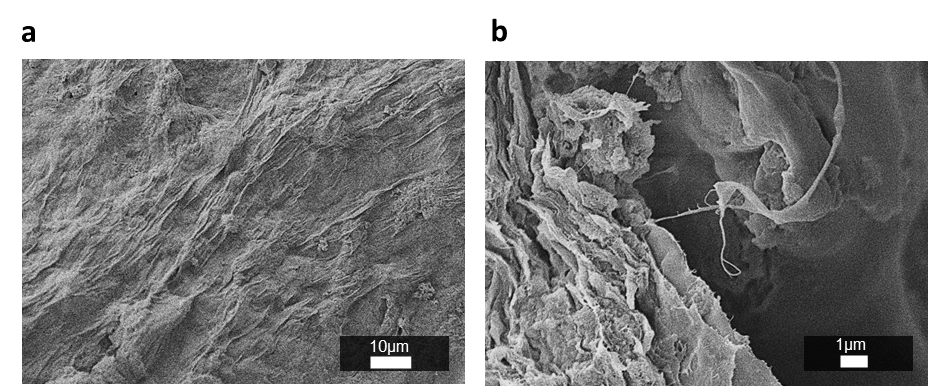

**Figure S12.** (a) Surface and (b) cross-sectional SEM images of DP_CNF_ film after mechanical testing.

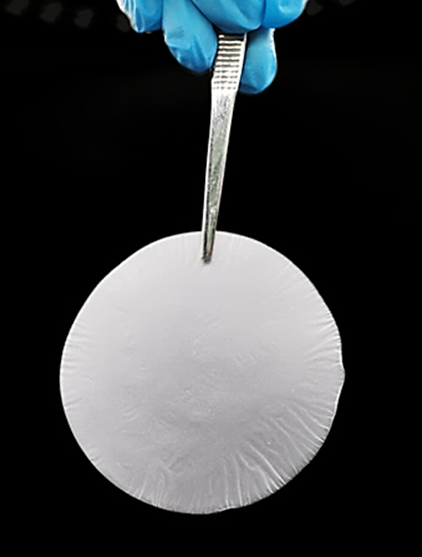

**Figure S13.** Wood_CNF_ nanopaper.

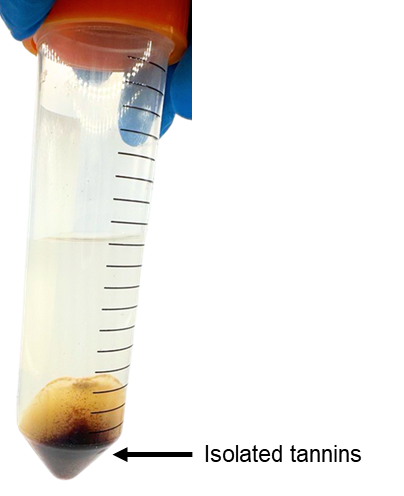

**Figure S14.** D_CNF_ upon centrifugation showing separation of tannins from CNF gel.

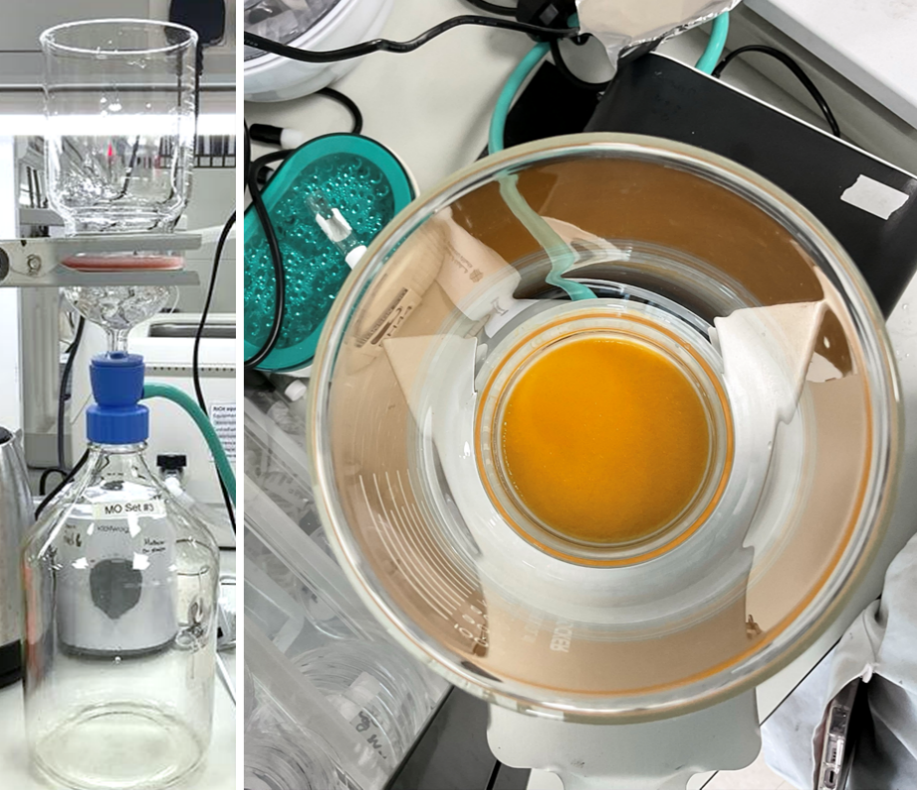

**Figure S15.** Vacuum filtration setup used to prepare the nanopapers.

Gwyddion AFM log:

file::nanoobserver(filename="C:\\Users\\CSI\\Documents\\CSI\\NanoSolution\\Malak\\251002 BWWMP 0.001% New protocol\\BWWMP 2 2025-10-02 16h25m56.nao")@2025-10-02 14:10:08.282452Z

proc::facet-level(mode=0)@2025-10-02 14:10:14.619923Z

proc::align_rows(direction=0, do_extract=False, do_plot=False, masking=0, max_degree=2, method=1, trim_fraction=0.05)@2025-10-02 14:10:18.664046Z

proc::fix_zero()@2025-10-02 14:10:20.163807Z

proc::fix_zero()@2025-10-02 14:10:21.820069Z

proc::scars_remove(max_width=4, min_len=16, threshold_high=0.666, threshold_low=0.25, type=5, update=True)@2025-10-02 14:10:22.932478Z

proc::scars_remove(max_width=4, min_len=16, threshold_high=0.666, threshold_low=0.25, type=5, update=True)@2025-10-02 14:10:23.683556Z

proc::scars_remove(max_width=4, min_len=16, threshold_high=0.666, threshold_low=0.25, type=5, update=True)@2025-10-02 14:10:24.395698Z

proc::scars_remove(max_width=4, min_len=16, threshold_high=0.666, threshold_low=0.25, type=5, update=True)@2025-10-02 14:10:24.747831Z

proc::scars_remove(max_width=4, min_len=16, threshold_high=0.666, threshold_low=0.25, type=5, update=True)@2025-10-02 14:10:24.932832Z

proc::scars_remove(max_width=4, min_len=16, threshold_high=0.666, threshold_low=0.25, type=5, update=True)@2025-10-02 14:10:25.102527Z

proc::scars_remove(max_width=4, min_len=16, threshold_high=0.666, threshold_low=0.25, type=5, update=True)@2025-10-02 14:10:25.267526Z

proc::scars_remove(max_width=4, min_len=16, threshold_high=0.666, threshold_low=0.25, type=5, update=True)@2025-10-02 14:10:25.443695Z

proc::scars_remove(max_width=4, min_len=16, threshold_high=0.666, threshold_low=0.25, type=5, update=True)@2025-10-02 14:10:25.611695Z

proc::facet-level(mode=0)@2025-10-02 14:10:26.817983Z

proc::fix_zero()@2025-10-02 14:10:30.931427Z

proc::align_rows(direction=0, do_extract=False, do_plot=False, masking=0, max_degree=2, method=1, trim_fraction=0.05)@2025-10-02 14:10:38.145541Z

proc::fix_zero()@2025-10-02 14:10:40.020207Z

proc::scars_remove(max_width=4, min_len=16, threshold_high=0.666, threshold_low=0.25, type=5, update=True)@2025-10-02 14:10:57.427225Z

proc::scars_remove(max_width=4, min_len=16, threshold_high=0.666, threshold_low=0.25, type=5, update=True)@2025-10-02 14:10:58.131800Z
